# Hydroxynonenal causes Langerhans cell degeneration in the pancreas of Japanese macaque monkeys

**DOI:** 10.1101/2021.01.07.425690

**Authors:** Piyakarn Boontem, Tetsumori Yamashima

## Abstract

**Background:** For their functions of insulin biosynthesis and glucose- and fatty acid- induced insulin secretion, the Langerhans β-cells require an intracellular milieu rich in oxygen. This requirement makes β-cells, with their constitutively low antioxidative defense, susceptible to the oxidative stress. Although much progress has been made in identifying its molecular basis in the experimental systems, whether the oxidative stress due to excessive fatty acids plays a crucial role in the Langerhans degeneration in primates is still debated.

**Methods:** Focusing on Hsp70.1, which has dual functions as a molecular chaperone and lysosomal stabilizer, the mechanism of lipotoxicity to the Langerhans islet cells was studied using Japanese macaque monkeys (*Macaca fuscata*) after the consecutive injections of the lipid peroxidation product hydroxynonenal. Based on the ‘calpain-cathepsin hypothesis’ of ischemic neuronal death formulated in 1998, calpain activation, Hsp70.1 cleavage, and lysosomal integrity were studied by immunofluorescence histochemistry, electron microscopy and Western blotting.

**Results:** Light microscopy showed higher vacuole formation in the treated islet cells than in the control cells. Electron microscopy showed that vacuolar changes that were identified as enlarged rough endoplasmic reticula occurred mainly in β-cells followed by δ-cells. Intriguingly, both cell types showed a marked decrease in insulin and somatostatin granules. Furthermore, they exhibited marked increases in peroxisomes, autophagosomes/autolysosomes, lysosomal and peroxisomal membrane rupture/permeabilization, and mitochondrial degeneration. Disrupted peroxisomes were often localized in the close vicinity of degenerating mitochondria or autolysosomes. Immunofluorescence histochemical analysis showed an increased colocalization of activated μ-calpain and Hsp70.1 with the extralysosomal release of cathepsin B. Western blotting showed increases in μ-calpain activation, Hsp70.1 cleavage, and hydroxynonenal receptor GPR109A expression.

**Conclusions:** Taken together, these data implicate hydroxynonenal in both the carbonylation of Hsp70.1 and the activation of μ-calpain. The calpain-mediated cleavage of the carbonyl group on Hsp70.1 after the hydroxynonenal-mediated carbonylation of Hsp70.1, may cause lysosomal membrane rupture/permeabilization. The low defense of primate Langerhans cells against exogenous hydroxynonenal and peroxisomally-generated hydrogen peroxide, was presumably overwhelmed to facilitate cell degeneration.

## Introduction

The intracellular milieu of pancreatic β-cells is rich in oxygen, glucose and fatty acids for insulin biosynthesis and secretion. This makes β-cells, with their constitutively low enzymatic antioxidative defense equipment [1,2], susceptible to oxidative stress during glucolipotoxicity. However, the mechanism of β-cell degeneration and death is not fully understood in type 2 diabetes until now. Hyperlipidemia is accepted to be the greatest risk factor, a phenomenon known as lipotoxicity [3], because non-esterified fatty acids (NEFAs) are potent inducers of reactive oxygen species (ROS) through different mechanisms. β-cells exposed to oxidative, excessive or conjugated NEFAs develop cell degeneration/death [3,4,5]. The molecular mechanisms underlying the oxidative injury due to ROS, comprise of endoplasmic reticulum (ER) stress, mitochondrial dysfunction, impaired autophagy, lysosomal disintegrity, inflammation, and mixture of them [6,7,8,9,10,11].

These molecular events are grossly similar to the neurodegeneration of hypothalamic arcuate nucleus occurring after the intake of high-fat diets [12] or hippocampal CA1 sector exposed to the oxidative stress during reperfusion after transient ischemia [3]. In 1998, Yamashima and his colleagues formulated the ‘calpain-cathepsin hypothesis’ as a mechanism of ischemic neuronal death [14,15]. Thereafter, they suggested role of lysosomal rupture due to calpain-mediated cleavage of oxidized (carbonylated) heat-shock protein 70.1 (Hsp70.1; human type of Hsp70, also called Hsp72) as a mechanism of neuronal death [5,16,17,18]. However, previous researchers have not found so far any convincing molecular mechanism of a major contribution of ROS to the β-cell degeneration and death. Antioxidative defence mechanisms of β-cells would be overwhelmed by overproduction of ROS [19] such as superoxide radicals (O_2_^•−^), hydrogen peroxide (H_2_O_2_), and the most reactive and toxic hydroxyl radicals (OH^•^). Among these, OH^•^ is responsible for the carbonylation of Hsp70.1 [16,17], and its precursor H_2_O_2_ plays a central role in the deterioration of glucose tolerance in the development of type 2 diabetes [19]. The source of H_2_O_2_ in β-cells has been considered to be the electron transport chain in mitochondria. While this source is obviously important, peroxisomal generation of H_2_O_2_ is more crucial in β-cells [20,21,22,23].

Lysosomes and peroxisomes, being contained in all eukaryote cells, are similar at the ultrastructural level but hold distinct enzymes for the completely different function. Lysosomes were discovered by the Belgian cytologist Christian de Duve in the 1950s [24]. They contain a wide variety of hydrolytic enzymes (acid hydrolases such as cathepsins B, L, etc.) which were produced in the rough ER, and are responsible for the digestion of macromolecules, old cell parts, and microorganisms. In oxygen-poor areas with an acidic environment (~pH4.5), lysosomes break down macromolecules such as damaged/aged proteins, nucleic acids, and polysaccharides for the recycling. For this reason, an ATP-driven proton pump is located inside the double membrane which borders the lysosomes in order to pump H^+^ ions into their lumen. The double membrane surrounding the lysosome is vital to ensure the cathepsins do not leak out into the cytoplasm and damage the cell from within. Recent data advocate for dual roles of Hsp70.1 not only as a molecular chaperone for damaged/aged/misfolded proteins but also as a guardian of lysosomal membrane integrity [25,26]. Therefore, in case of Hsp70.1 dysfunction, not only failure of protein traffic and degradation (autophagy failure) but also lysosomal destabilization (cell degeneration and death) may occur.

Peroxisomes are single membrane-bounded organelles that are best-known for their involvement in both the cellular lipid metabolism and the cellular redox balance [27,28,29]. Peroxisomes were first described by a Swedish doctoral student, J. Rhodin in 1954 [30], and were identified as a cell organelle by de Duve. He named them “peroxisomes”, replacing the formerly used morphological term “microbodies” [31]. De Duve and Baudhuin discovered that peroxisomes contain several oxidases involved in the production of H_2_O_2_ as well as catalase involved in the decomposition of H_2_O_2_ to oxygen and water [31,32]. So, ‘peroxisomes’ owe their name to ‘hydrogen peroxide’ -generating and -scavenging activities. Peroxisomes are small spheroid organelles with a fine, granular matrix. Peroxisomes hold diverse oxidative enzymes that were produced at free ribosomes and require oxygen. To absorb nutrients that the cell has acquired, peroxisomes digest long-chain fatty acids and break them down into smaller molecules by β-oxidation. One of the byproducts of the β-oxidation is H_2_O_2_, so peroxisomes break H_2_O_2_, down into water (H_2_O) and O_2_. H_2_O_2_ itself is not very reactive, but Fe^2+^ or Fe^3+^ reacts with H_2_O_2_ to produce OH^•^ in mitochondria. Accordingly, in both health and diseases, peroxisomes and mitochondria have a close relationship each other. Peroxisome generation, maintenance, and turnover are of great importance for the cellular homeostasis and survival. Peroxisomes are degraded by themselves or when fusing with lysosomes. The degradation takes just ~4 min, and half-life of peroxisomes is only 5 days.

Hydroxynonenal is a major aldehyde which was produced during the OH^•^-mediated peroxidation of ω-6 polyunsaturated fatty acids. In 2009, Oikawa et al. (2009) [16] found by the proteomic analysis that Hsp70.1 can be its target of carbonylation. Subsequently, Sahara and Yamashima (2010) [33] demonstrated by the in-vitro experiment that Hsp70.1 being carbonylated by hydroxynonenal is prone to the activated μ-calpain-mediated cleavage. Calpain-mediated cleavage of carbonylated Hsp70.1 leads to the lysosomal membrane rupture/permeabilization with the resultant release of cathepsins. Although much progress has been made in identifying the molecular basis of lipotoxicity in the experimental systems, whether or not this phenomenon actually plays an important role for the Langerhans cell degeneration remains debated especially in the primates [10]. To overcome this controversy, the authors thought that the experimental paradigm using non-human primates is indispensable. So, in the preliminarly study we confirmed that the intravenous injections of the synthetic hydroxynonenal induced cell death of diverse brain neurons and hepatocytes of the Japanese macaque monkeys [18]. Here, we focused on the pancreas of monkeys after the hydroxynonenal injections to elucidate the molecular cascade processing in the primate organ. The pancreas of monkeys is large enough to supply sufficient amount of tissues necessary for the simultaneous analyses of immunofluorescence histochemistry, light and electron microscopy, and Western blotting in the given animal.

## Materials and methods

### Animals

After the referee of animal experimentation about the ethical or animal welfare, four young (4~5 years: compatible with teenagers in humans) female Japanese macaque monkeys (*Macaca fuscata*) were supplied by National Bio-Resource Project (NBRP) “Japanese monkey” (National Institute for Physiological Sciences, Okazaki, Japan). After arrival, the monkeys were reared in the wide cage with autofeeding and autodrainage machines as well as appropriate toys to play at least for 1 year to facilitate acclimation.

The room temperature was kept 22~24 °C with the humidity of 40~50%. They were fed by 350 kCal/Kg body weight of non-purified solid monkey foods per day containing vitamines. and apples, pumpkins or sweet potatoes were given twice every week. In the morning and afternoon, an animal care staff and the first author monitored the health and well-being of the animals to check the consumption of foods, pupilar reflex to the light, and conditions of standing and jumping.

At 5~6 years of age, monkeys with body weight 5~7 Kg were randomly divided into two different groups of the sham-operated control (n=1) and those undergoing hyroxynonenal injections (n=3). In 3 monkeys, under the intramuscular anesthesia using 2 mg/Kg of kethamine hydrochrolide, intravenous injections of 5 mg/week of synthetic hydroxynonenal (Cayman Chemical, Michigan, USA) were done for 24 weeks. Such doses and serial injections were designed to temporarily mimic blood concentrations of hydroxynonenal in humans around 60’s [34]. Behavioural changes such as reduced exploration, standing, and jumping as well as decreases of appetite and body weight were carefully monitored to implement humane endpoints. However, all the monkeys were fine until the end of final 24th hydroxynonenal injections.

### Tissue collection

Six months after the initial injection and within a couple of weeks after the final injection, the monkeys were immobilized by the intramuscular injection of 10 mg/Kg BW ketamine hydrochloride followed by the intravenous injection of 50 mg/Kg BW sodium pentobarbital. In addition, to ameliorate animal suffering, the monkey was deeply anesthetized with 1.5% halothane plus 60% nitrous oxide. After the perfusion of 500 mL saline through the left ventricle, the pancreas was removed without suffering any pain. Half of the tissue was fixed in either i) 4% paraformaldehyde for light microscopy, or ii) 2.5% glutaraldehyde for electron microscopy. The remaining half was stocked in the −80 °C deep freezer for the Western blotting analysis.

### Histological and immunofluorescence histochemical analyses

The pancreas tissues after fixation with 4% paraformaldehyde for 2 weeks were embedded in paraffin, and 5μm sections were stained by hematoxylin-eosin. For the immunofluorescence histochemistry, the cryoprotected pancreas tissues embedded in the OCT medium (Sakura Finetek, Japan) were cut by cryotome (Tissue-Tek® Polar®, Sakura, Japan), and 5μm sections were immersed with heated 0.01% Citrate retrieval buffer to induce epitope retrieval. Non-specific staining was blocked with 1% bovine serum albumin (Nacalai tesque, Japan), and were incubated overnight at 4°C with the primary antibodies at the dilution of 1:100. We used mouse monoclonal anti-human Hsp70 (BD Bioscience, USA), rabbit anti-human activated μ-calpain (order made by PEPTIDE Institute, Japan), rabbit anti-human cathepsin B (Cell Signaling, USA), and mouse monoclonal anti-Lamp2 (Abcam, USA) antibodies. After washings, the sections were incubated for 30 min with secondary antibodies; Alexa Fluor™ 594 goat anti-mouse IgG [H+L] (Invitrogen, USA), or Alexa Fluor™ 488 goat anti-rabbit IgG (Invitrogen, USA) at the dilution of 1:500. To block autofluorescent staining, Autofluorescence Quenching Kit (Vector Laboratories, USA) was utilized. The immunoreactivity was observed with the laser confocal microscope (LSM5 PASCAL, Software ZEN 2009, Carls Zeiss, Germany).

### Ultrastructural analyses

For the electron microscopic analysis, small specimens of the pancreas tissue were fixed with 2.5% glutaraldehyde for 2h and 1% OsO_4_ for 1h. Subsequently, they were dehydrated with graded acetone, embedded in resin (Quetol 812, Nisshin EM Co. Tokyo), and thin sections were made. After trimming with 0.5% toluidine blue-stained sections, the ultrathin (70nm) sections of appropriate portions were stained with uranyl acetate (15 min) and lead citrate (3 min), and were observed by the electron microscope (JEM-1400 Plus, JEOL Ltd., Tokyo).

### Western blotting

Total protein extraction was done, using protease inhibitor cocktail (Sigma-Aldrich, USA) and PhosSTOP phosphatase inhibitor cocktails tablets (Roche, Germany). After centrifugation at 12,000 rpm for 10 min, the supernatant proteins were determined by Bradford Assay (Thermo Fisher, USA). Twenty μg proteins were separated by SDS-PAGE in SuperSep (TM) Ace 5-20% gel (Wako, Japan) at 40 mA for 1h. The total proteins were transferred to PVDF membrane (Millipore, USA). Transferred protein quantities were detected with Ponseau S solution (Sigma-Aldrich, USA). Transferred proteins were blocked with 1% BSA (KPL Detector™ Block, USA) for 1h. The blots were incubated with mouse monoclonal anti-human Hsp70 antibody (BD Bioscience, USA) at the dilution of 1:4,000, rabbit anti-human activated μ-calpain antibody (PEPTIDE Institute, Japan) at 1:250 overnight, or rabbit anti-human GPR109A (also called hydroxycarboxylic acid receptor 2) antibody (Abcam, USA) at 1:500. β-actin was utilized as an internal control at a dilution of 10,000 (Sigma-Aldrich, USA). The immunoblots were subsequently incubated for 1h with secondary antibodies at 1:10,000 dilution of anti-mouse (Santa Cruz, USA) or anti-rabbit IgG (Sigma, USA). An enhanced chemiluminescence (ECL) HRP substrate detection kit (Millipore, USA) was used to visualize the reactive protein bands with ImageQuant LAS 4000 mini (GE Life Science, USA).

### Ethics

This study was carried out in strict accordance with the recommendations in the Guide for the Care and Use of Laboratory Animals of the National Institutes of Health. The protocol was approved by the Committee on the Ethics of Animal Experiments of the Kanazawa University Graduate School of Medical Sciences (Protocol Number: AP-153613).

## Results

By light microscopy, the Langerhans islet cells after the hydroxynonenal injections showed formation of many vacuoles, compared to the control, which were often filled with eosinophilic substance or looked almost empty (Fig. 1). Nuclear chromatin was generally more dense than the control, which was consistent with the electron microscopic finding (Fig. 2). A small number of nuclei showed dissolution of chromatin or punctuate condensation (Fig. 1, dot circle). However, neither apoptotic bodies nor membrane blebbings were observed in the Langerhans islet cells after the hydroxynonenal injections.

**Figure 1.**
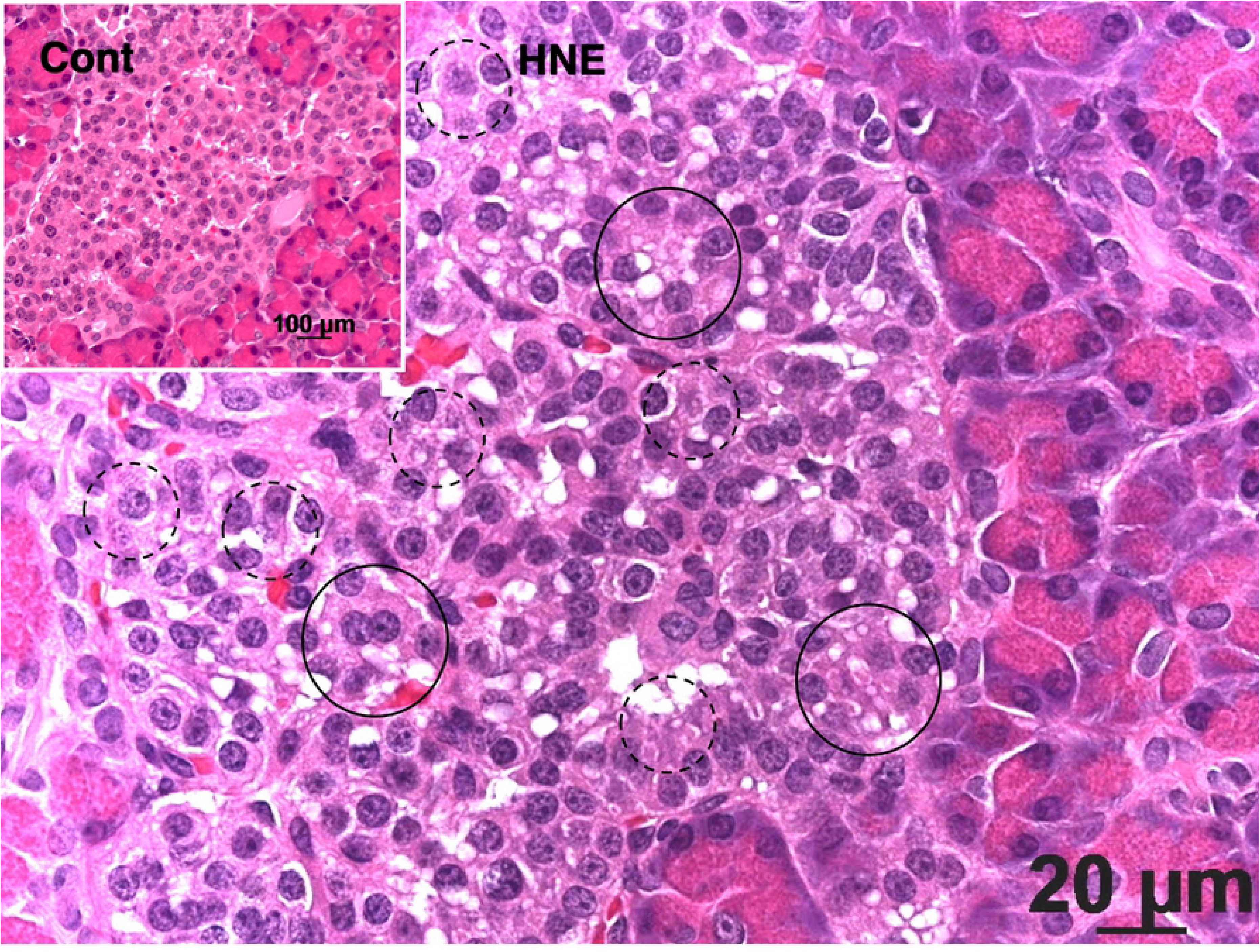
Light micrograph of the monkey pancreas. The Langerhans islet after the hydroxynonenal injections (HNE) shows many vacuole formation (circles), compared to the sham-operated control (Cont). Nuclear chromatin is generally more dense after the hydroxynonenal injections, compared to the control. A small number of nucleus shows diffuse dissolution or punctuate condensation (dot circle). However, neither apoptotic bodies nor membrane blebbings were seen. Acinar cells are distributed in the surrounding area. Hematoxylin-eosin staining.

**Figure 2.**
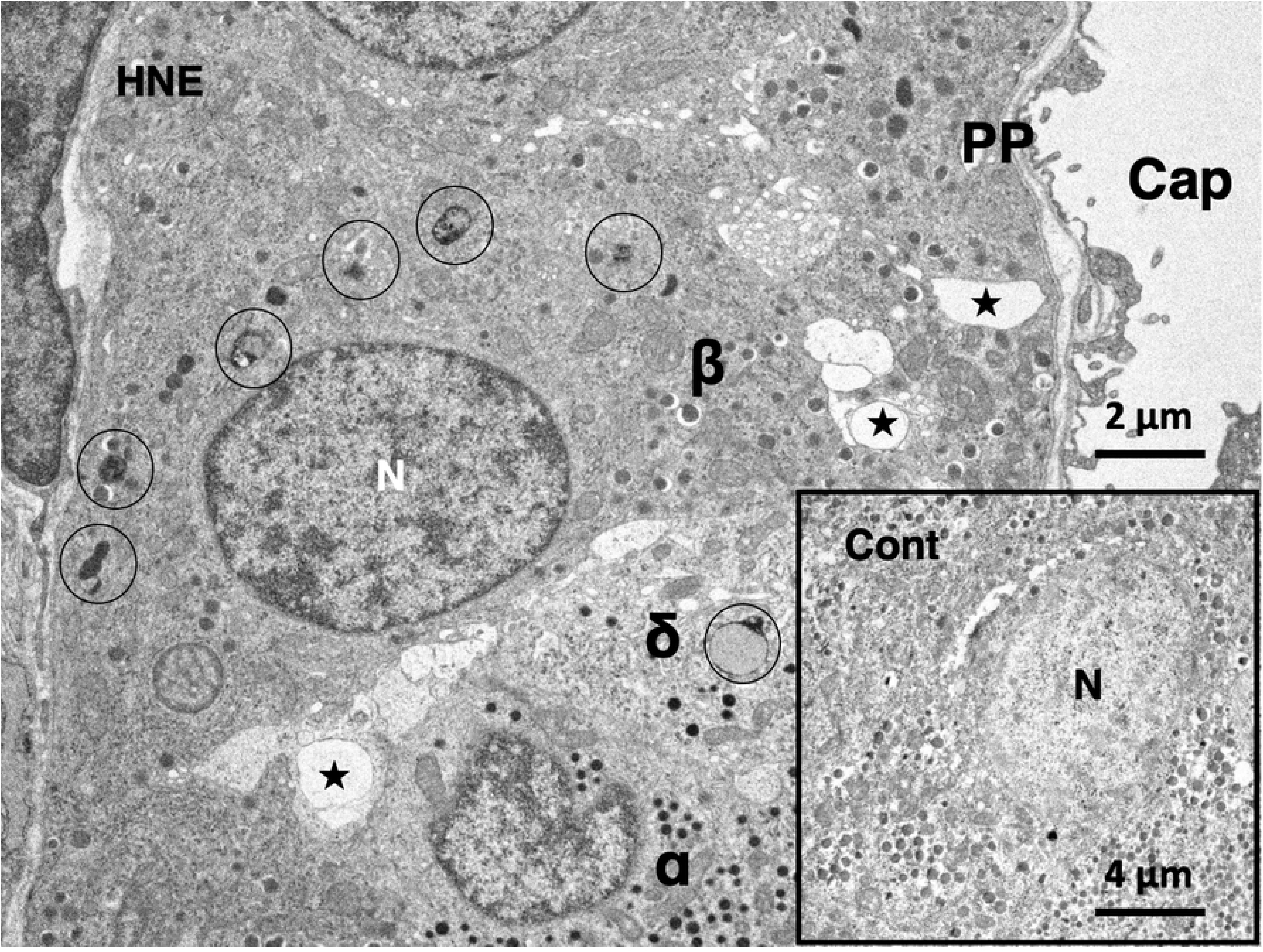
Electron micrograph of the Langerhans islet of the monkey before and after the hydroxynonenal injections. The Langerhans islet cells are situated surrounding a capillary (Cap), and each cell is characterized by peculiar secretory granules. Insulin secretory granules of the β-cell (β) are the largest with a clear halo. Glucagon secretory granules of the α-cell (α) lack a clear halo, being a little bit smaller but more electron-dense than insulin granules. Somatostatin-containing granules of the δ-cell (δ) are also electron-dense like glucagon granules, showing a sparse distribution. PP-cells (PP) contain spherical or elliptical small granules, which are very heterogeneous in size. The greatest changes after the hydroxynonenal injections are a remarkable decrease of insulin and somatostatin granules as well as an increase of autophagosomes and autolysosomes. In the control β-cell (Cont), insulin granules were distributed throughout the cytoplasm. In contrast, in the β-cell after the hydroxynonenal injections (HNE), the distribution of insulin granules was restricted in the cytoplasm toward the capillary. Instead, especially β-cell showed formation of many autolysosomes (circles), compared to the control (Cont). Further, both β-cell and δ-cell showed vacuole formations (stars) and more electron-dense nuclear chromatin. N; nucleus

The Langerhans islets are known to be basically composed of α-, β-, δ- cells and pancreatic polypeptide cells (PP-cells). As described previously in humans [35,36], electron microscope was useful for differentiating 4 types of cells in the Langerhans islets of monkeys. In β-cells (β), insulin secretory granules had an electron-opaque core of 300-400 nm with a clear halo (Figs. 2–5), while in α-cells (α), glucagon secretory granules were electron-dense without a clear halo. Compared to the insulin secretory granules, the glucagon granules were more electron-dense with a smaller diameter (200-300 nm) (Fig. 2). δ-cells (δ) exhibit neuron- or trumpet- like morphology with cytoplasmic processes extending from the islet capillaries (Fig. 2). Somatostatin-containing granules were also electron-dense, showing a similar size with glucagon granules, but showed a sparse distribution (Fig. 6a,b). PP-cells (PP) contained spherical smaller granules, which were heterogeneous in size (Fig. 2).

By the electron microscopic observation, the most remarkable change in the Langerhans islets after the hydroxynonenal injections was a remarkable decrease of insulin and somatostatin granules (Fig. 2, HNE), compared to the control (Fig. 2, Cont). The decrease was evident, when compared with human β- and δ- cells whose cytoplasm was filled with insulin and somatostatin granules [36]. In the control β-cells, insulin granules were distributed throughout the cytoplasm (Fig. 2, Cont), whereas in the Langerhans islets after the hydroxynonenal injections, they were clustered in the cytoplasm toward the capillary lumen (Fig. 2, HNE). Among α-, β-, δ- and PP- cells of the islets after the hydroxynonenal injections, β- and δ- cells showed lots of vacuole formations which were revealed to be enlarged rough ER with a marked decrease of ribosomes (Fig. 2 stars, Fig. 5), indicating ER dysfunction. Further, β- and δ- cells after the hydroxynonenal injections were characterized by peroxisomal proliferation (Figs. 5, 6b), and increments of autophagosomes or autolysosomes (Figs. 2–5, 6a,b), which were extremely rare in the control tissue (Fig. 2, Cont).

By the electron microscopic observation of the pancreas, lysosomes were relatively electron-dense organelles homogenously filled with microvesicular granules, while peroxisomes contained a less-dense matrix with fine granules. So, the limiting membrane was more clearly visible in peroxisomes than lysosomes. In the β-cells after the hydroxynonenal injections, lysosomes were often distributed around the autophagosomes or autolysosomes throughout the cytoplasm (Figs. 2, 3), while peroxisomes were distributed around insulin granules, being intermingled with degenerating mitochondria in the cytoplasm toward the capillary lumen (Fig. 5). At the high magnification, degenerating β-cells (Fig. 3) and δ-cells (Fig. 6a) contained an autophagosome which was in the process of fusing with the lysosome showing membrane permeabilization. Autophagosome or autolysosome was observed to fuse also with peroxisome which was surrounded by the disrupted membrane (Figs. 4, 6b). Peroxisomes with membrane disruption were abundant around insulin and somatostatin granules in the cytoplasm toward the capillary lumen (Fig. 4, 6a). Some peroxisomes were seen to fuse with degenerating mitochondria (Fig. 5, circles) or in the close vicinity of autolysosomes (Fig. 5, dot circles). As β-cells and δ-cells showed a similar change, β-cells generally showed a stronger degeneration than δ-cells.

**Figure 3.**
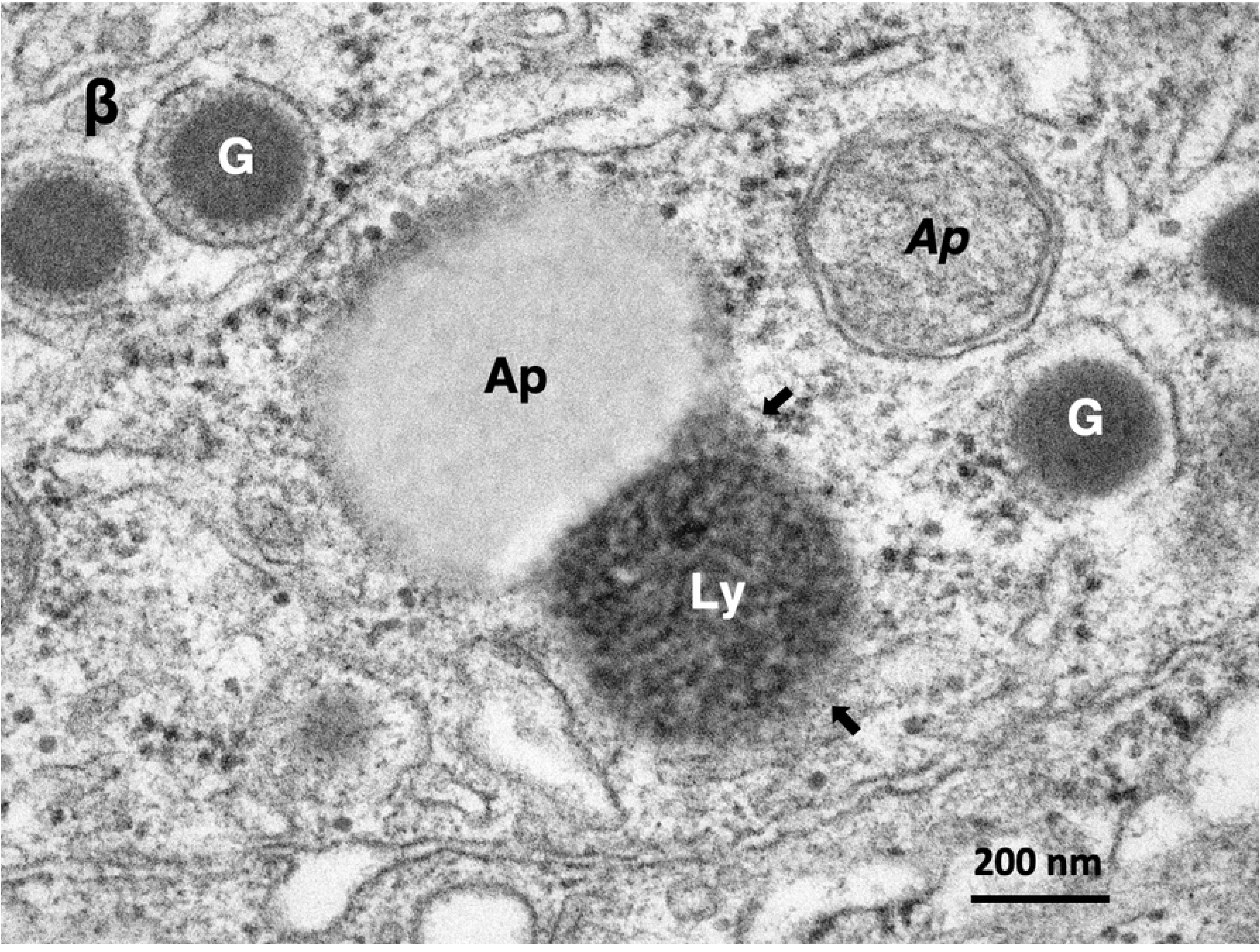
Electron micrograph of the β-cell in the monkey after the hydroxynonenal injections. β-cell shows fusion of an autophagosome (AP) with a lysosome (Ly) exhibiting evidence of membrane permeabilization (arrows). This autophagosome is devoid of double membrane, whereas another one (*Ap*) presumably containing degenerated mitochondria, has distinct double membrane. G: insulin granule,

**Figure 4.**
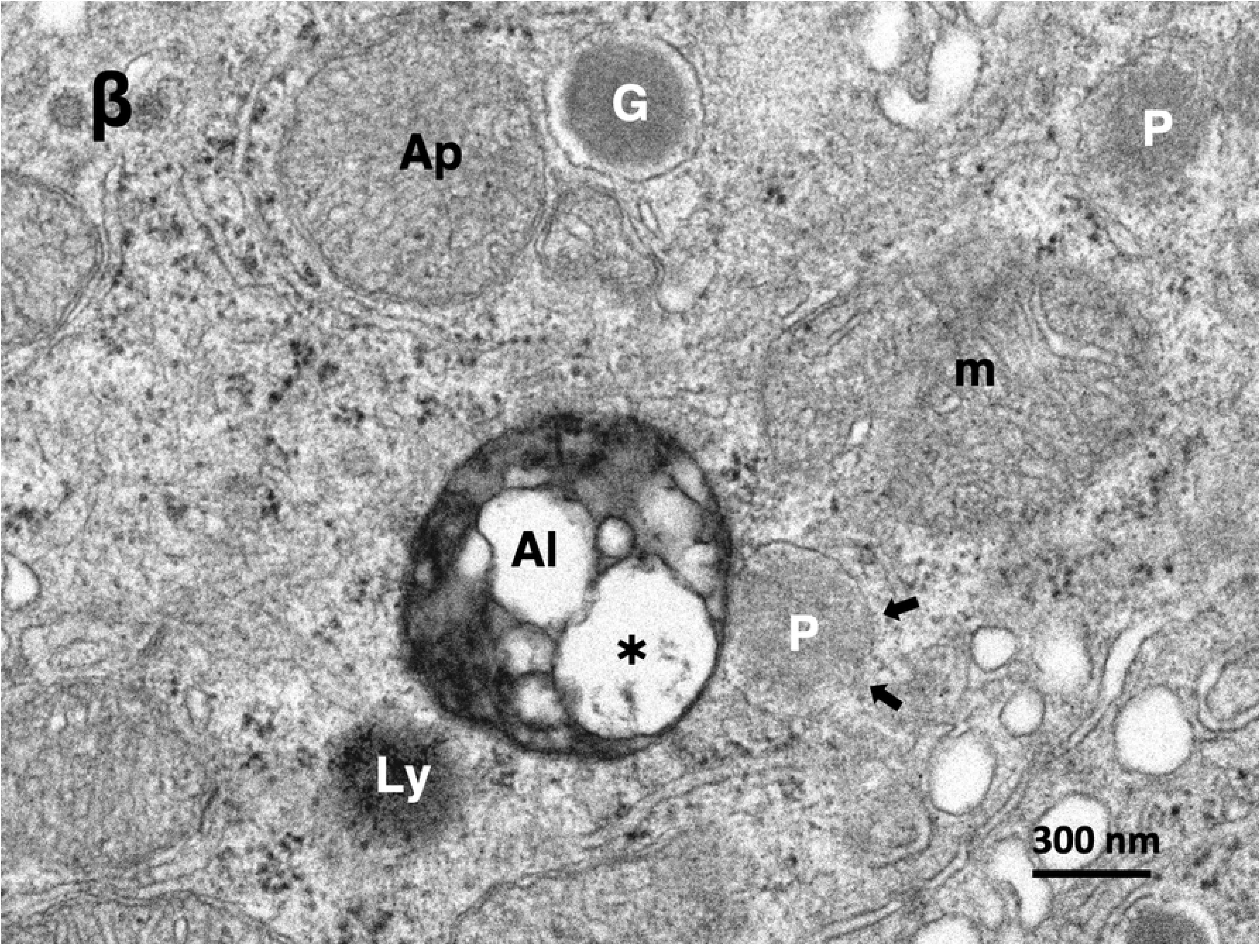
Electron micrograph of the β-cell in the monkey after the hydroxynonenal injections. β-cell shows fusion of an autolysosome (Al) with a peroxisome (P) exhibiting membrane disruption (arrows). The autolysosome (Al) contains mitochondria-derived debris (asterisk). The neighbouring lysosome (Ly) shows membrane permeabilization. G: insulin granule, Ap: autophagosome, m: degenerating mitochondria

**Figure 5.**
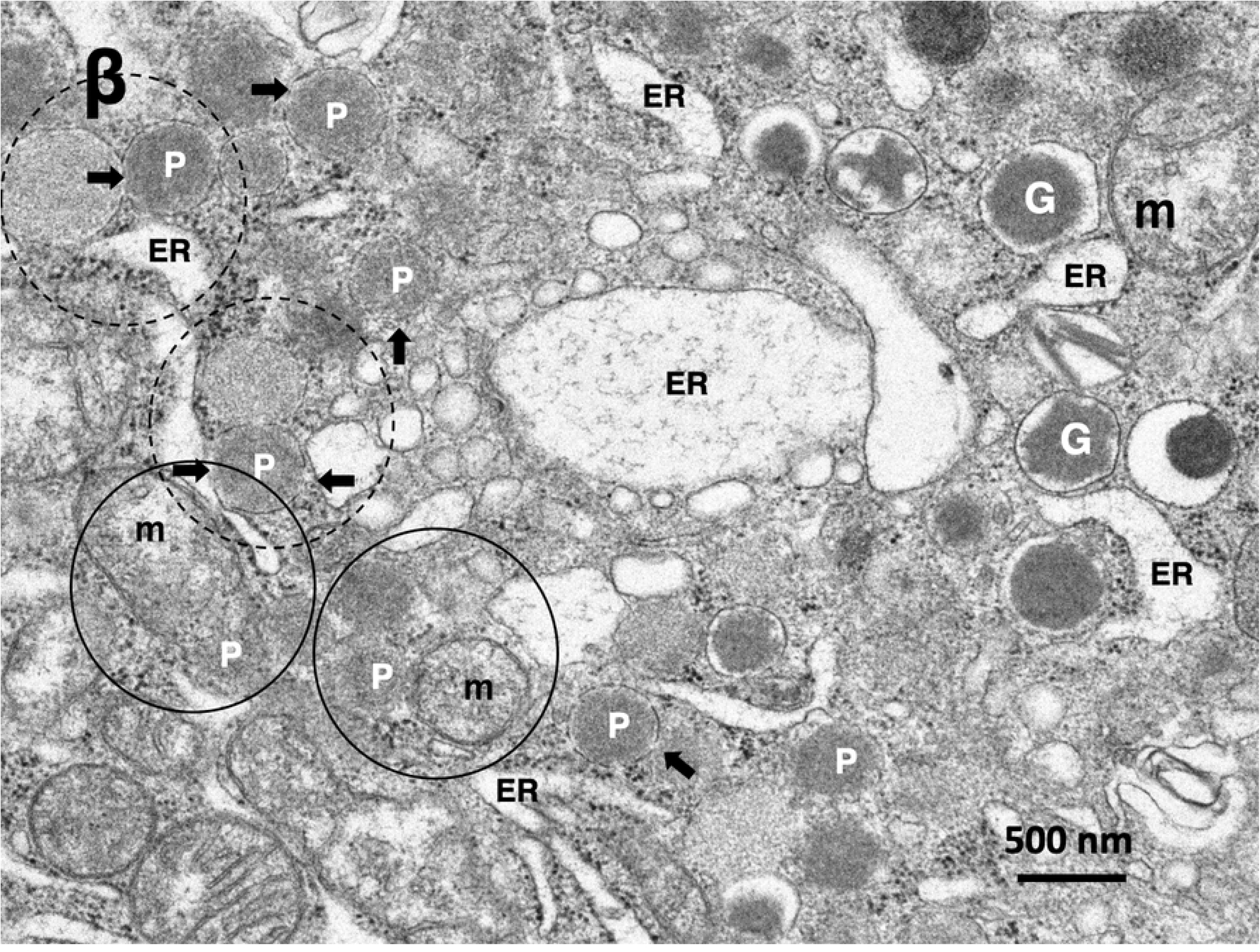
Electron micrograph of the β-cell in the monkey after the hydroxynonenal injections. β-cell shows enlargement of rough ER (ER), as seen in Fig. 2. Proliferation of peroxisomes (P) is obvious, and they are intermingled with degenerating mitochondria and insulin granules in the cytoplasm toward the capillary lumen, as seen in Fig. 2. Some peroxisomes with membrane disruptions (arrows) are fusing with degenerating mitochondria (m, circles) or autolysosomes presumably containing mitochondrial debris (dot circles). G: insulin granule

**Figure 6.**
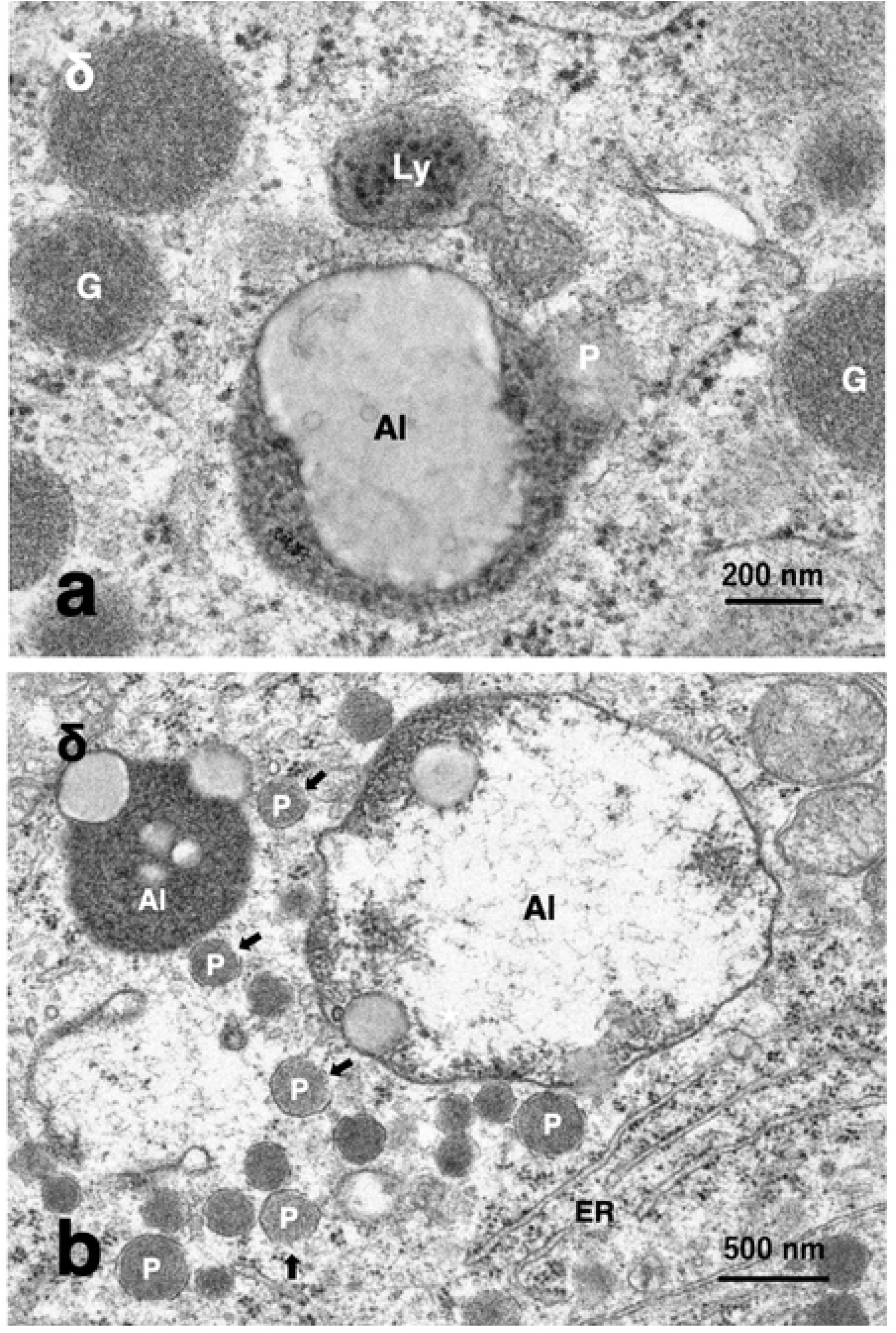
**6a) Electron micrograph of the δ-cell in the monkey after the hydroxynonenal injections. δ-cell** shows fusion of an autolysosome (Al) with a peroxisome (P) exhibiting membrane disruption. G: somatostatin granule without a halo, Ly: lysosome **6b) Electron micrograph of the δ-cell in the monkey after the hydroxynonenal injections.** δ-cell shows peroxisomal proliferation (P) around the autolysosomes (Al). The latter are presumably in the process of fusing with peroxisomes (P). Some peroxisomes show membrane disruption (arrows). ER: rough ER

Immunofluorescence histochemical analysis showed a remarkable increase of Hsp70.1 colocalization with activated μ-calpain in the Langerhans islets after the hydroxynonenal injections (Fig. 7a, HNE, merged color), comparted to the control (Fig. 7a, Cont). In the control, the cathepsin B immunoreactivity was seen as tiny granules which were compatible with the size of intact lysosomes (Fig. 7b, Cont). However, after the hydroxynonenal injections, cathepsin B immunoreactivity was seen as coarse granules with a diffuse, thin distribution throughout the cytoplasm (Fig. 7b, HNE). In the core of cathepsin B immunoreactivity, lamp-2 immunoreactivity was colocalized with a merged color (data not shown here), which showed extralysosomal release of cathepsin enzymes by the lysosomal membrane rupture/permeabilization.

**Figure 7.**
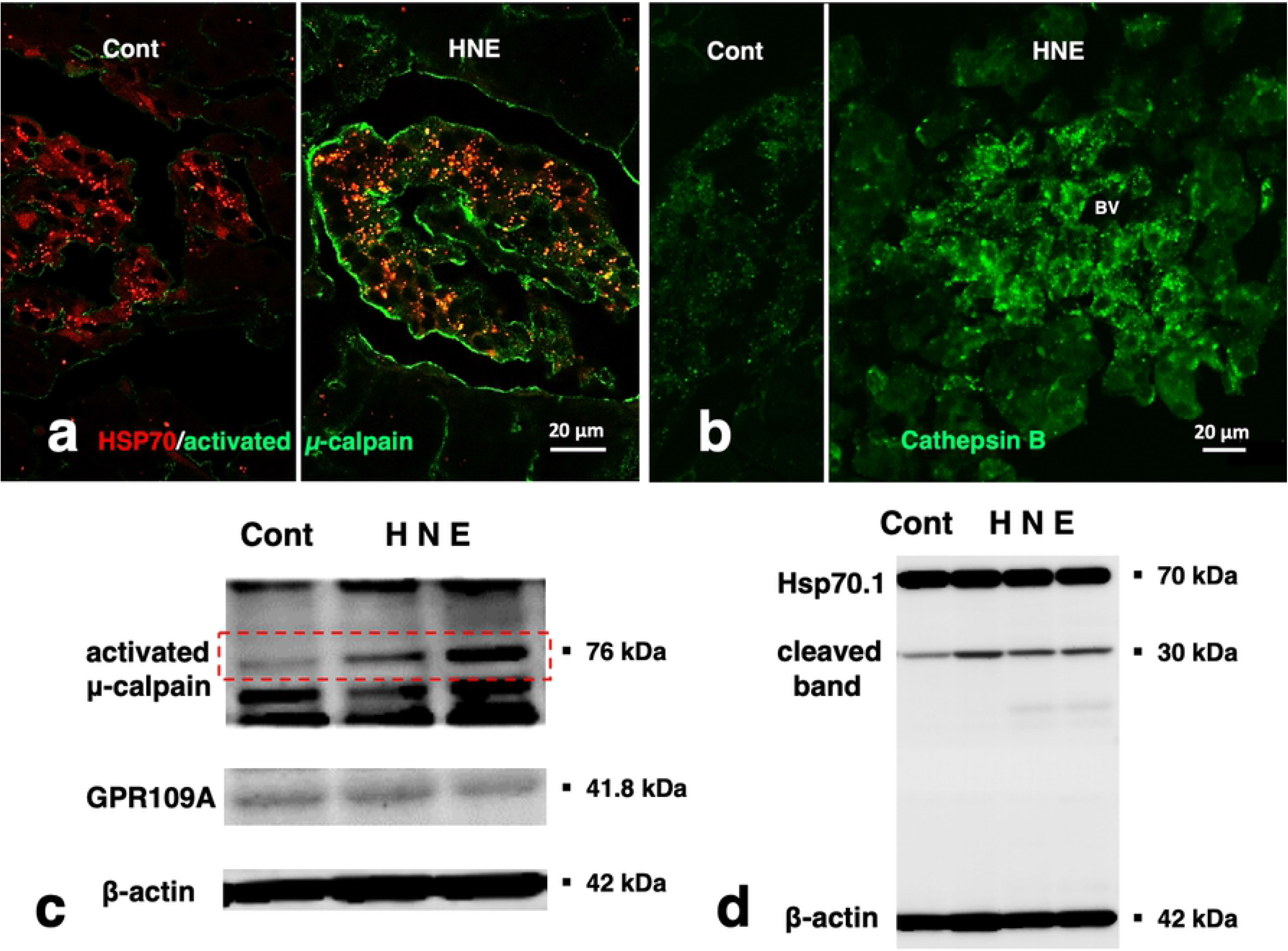
Calpain activation, Hsp70.1 cleavage, and cathepsin B leakage. **7a) Immunofluorescence histochemical staining of Hsp70.1 (red) and activated μ-calpain (green).** Activated μ-calpain immunoreactivity is negligible without hydroxynonenal injections (Cont), whereas μ-calpain activation occurs with hydroxynonenal injections (HNE), being consistent with the Western blotting data (c, activated μ-calpain). After hydroxynonenal injections, activated μ-calpain immunoreactivity (green) is colocalized with Hsp70.1 immunoreactivity (red), showing a merged color of yellow (HNE, yellow). The distribution pattern of granular merged colors is compatible with that of autolysosomes as seen in Fig. 2. **7b) Immunofluorescence histochemical staining of cathepsin B (green).** Cathepsin B is stained as tiny granules in the control Langerhans islet (Cont), whereas stained as coarse granules with the perigranular immunoreactivity after hydroxynonenal injections (HNE), which indicates lysosomal membrane rupture/permeabilization. The nuclei show negligible cathepsin B immunoreactivity. Cathepsin B immunoreactivity was colocalized with that of Lamp2 (data not shown here). **7c, d) Western blotting analyses of activated μ-calpain (c, 76 kDa), GPR109A (c, 41.8 kDa), and Hsp70.1 (d, 70 kDa).** Compared to the control (Cont), μ-calpain is activated after hydroxynonenal injections (c, red rectangle), while Hsp70.1 cleavage is increased after hydroxynonenal injections in all three monkeys (d, HNE), compared to the control (Cont). GPR109A is expressed in the pancreas tissue, although showing no change after hydroxynonenal injections (c, GPR109A).

Western blotting showed an increase of 76 kDa band intensities of activated μ-calpain after the hydroxynonenal injections, compared to the control (Fig. 7c, red rectangle). Compared to the control, calpain-mediated Hsp70.1 cleavage [33] increased in all three monkeys after the hydroxynonenal injections (Fig. 7d). Repeated Western blotting showed the same results, although the statistical significance was not available because of the restricted number of experimental animals. GPR109A was expressed equally in the control and hydroxynonenal-injected tissues (Fig. 7**c**).

## Discussion

In type 2 diabetes, β-cell degeneration and death contribute to the loss of its functional mass [8,37,38]. However, surprisingly, the underlying molecular mechanism remains grossly unknown until now, although β-cell lipotoxicity has been subject to intensive research for the past two decades [39,40,41]. As most studies showing deleterious effects of elevated NFFAs on β-cell function were conducted *in vitro,* the contribution of elevated NEFAs to β-cell dysfunction in human patients with type 2 diabetes still remains controversial [41]. Not only quantity but also quality (i.e. extent and type of oxidization) of NEFAs may impact β-cell function [10].

In the pancreas, two types of fatty acid receptor emerge as a cause of abnormal Ca^2+^ mobilization in response to excessive and/or oxidized NEFAs; one is G protein-coupled receptor 40 (GPR40) while another is GPR109A. GPR40 is a receptor for diverse NEFAs, being expressed abundantly in the β-cells [42,43]. Steneberg et al. (2005) [44] proposed that GPR40 is indirectly responsible for hyperinsulinemia-induced insulin resistance, whereas other researchers reported that NEFA-induced hyperinsulinemia represents a mechanism by which β-cells attempt to compensate for the insulin resistance because this ability is compromised by GPR40 deletion [45,46]. Accordingly, implication of GPR40 for the development of β-cell lipotoxicity in response to excessive NEFAs still remains debated [5]. Albeit GPR40 is abundantly expressed in the primate pancreas [42,43] including the present macaque monkeys (data not shown), for this reason we focused here on the expression of a metabolic sensor GPR109A in the monkey pancreas. Since in response to hydroxynonenal GPR109A was demonstrated to induce excessive Ca^2+^ mobilization and the subsequent cell death in the retinal pigmented and colon epithelial cells [47], we speculated that the same may occur in β-cells where GPR109A is expressed.

Previous reports suggest that oxidative stress during metabolizing excessive NEFAs mediates lipotoxicity. As NEFAs are metabolized not only through mitochondrial β-oxidation but also through peroxisomal β-oxidation, the subcellular sites of ROS formation are mitochondria and peroxisomes [27,28,48]. In contrast to the mitochondrial β-oxidation, however, the acyl-CoA oxidases generate more H_2_O_2_ in the peroxisomes [49]. Now, almost 50 years after their discovery, it is well known that peroxisomes can function as a main source of ROS [50]. As β-cells almost completely lack catalase, they are thought to be exceptionally vulnerable to abundant H_2_O_2_ which was presumably generated in the increased number of peroxisomes (Fig. 5).

It is well known that peroxisomes are highly plastic, dynamic organelles that rapidly modulate their size, number, and enzyme content in response to changing environmental conditions [51,52]. The present study showed a remarkable peroxisomal proliferation in β-cells after the hydroxynonenal injections (Figs. 5, 6b). Inactivation of H_2_O_2_ by catalase seems to be a step of critical importance for the removal of ROS in β-cells. However, since β-cells almost completely lack the H_2_O_2_-detoxifying enzyme, oxidoreductase catalase [1,2], they are exceptionally vulnerable to H_2_O_2_ that was generated in peroxisomes. If H_2_O_2_ is not quickly converted into water and oxygen by catalase, it can react in an iron-catalysed reaction with O_2_^•−^ yielding the highly reactive OH^•^. It is the most reactive oxygen radical known, reacting instantaneously with molecules in its immediate vicinity, which explains its great destructive power. This conceivably causes β-cell dysfunction, degeneration, and ultimately cell death, because of its low antioxidative defense status [23]. Since mitochondria also lack catalase and, in β-cells, mitochondria contain an extremely low glutathione peroxidase activity, any excessive oxidative stress is detrimental for the β-cell mitochondria due to the limited antioxidative defence capacity [19]. The capacity for inactivation of H_2_O_2_ is sufficient under normal circumstances, however, during the consecutive oxidative stress like the present experimental paradigm (5mg/W x 24 injections of hydroxynonenal), a constant level of H_2_O_2_ might have been continuously generated within both mitochondria and peroxisomes. So, low anti-oxidative defence of β-cells can be easily overwhelmed in pathological situations of sustained OH^•^ production. The close spatial relation of mitochondria with crista disruption and peroxisome with membrane disruption (Fig. 5), indicates a strong impact of hydroxynonenal upon β-cells.

Peroxisomal degradation occurs through at least three different mechanisms: macropexophagy, micropexophagy, and 15-lipoxygenase-mediated autolysis [53,54]. A lipid-peroxidizing enzyme, 15-lipoxygenase can generate hydroxynonenal from ω-6 polyunsaturated fatty acids of biomembranes. As 15-lipoxygenase is distributed in peroxisomes at the highest concentration [55], it initiates organelle degradation by pore formation in the limiting membranes [56,57]. van Leyen et al. (1998) [58] demonstrated that 15-lipoxygenase can bind to organelle membranes and induce the leakage of its contents. Yokota et al. (2001) [55] also showed induction of the peroxisomal membrane disruption and the resultant catalase leakage by 15-lipoxygenase in the rat liver. Interestingly, such 15-lipoxygenase-induced peroxisomal membrane disruptions were very similar at the ultrastructural level to those observed in the monkey Langerhans islet cells after the hydroxynonenal injections (Figs. 5, 6b). Hydroxynonenal is generated as a consequence of H_2_O_2_-mediated lipid peroxidation of biomembranes composing of ω-6 polyunsaturated fatty acids such as linoleic and arachidonic acid [11,59,60]. Therefore, it is reasonable to speculate that exogenous hydroxynonenal is a causative substance and also a trigger, which induce consecutive peroxisomal HO^•^ production with the resultant intrinsic hydroxynonenal generation in the monkey pancreas (Fig. 8). Hydroxynonenal has been increasingly recognized as a particularly important mediator of dysfunction and degeneration of pancreatic β cells in type 2 diabetes and alcohol-induced pancreatic damage [19,61]. There is ample evidence that hydroxynonenal causes cell dysfunction and death in diverse disorders including Alzheimer’s disease [62,63], cardiovascular disease, stroke, arthritis and asthma [64,65,66,67]. By activating GPR109A, hydroxynonenal can be a trigger for the excessive Ca^2+^ mobilization and subsequent calpain activation [68]. So, it is reasonable that the Langerhans islet cells with GPR109A expression showed an increased activation of μ-calpain in response to the exogenous hydroxynonenal.

**Figure 8.**
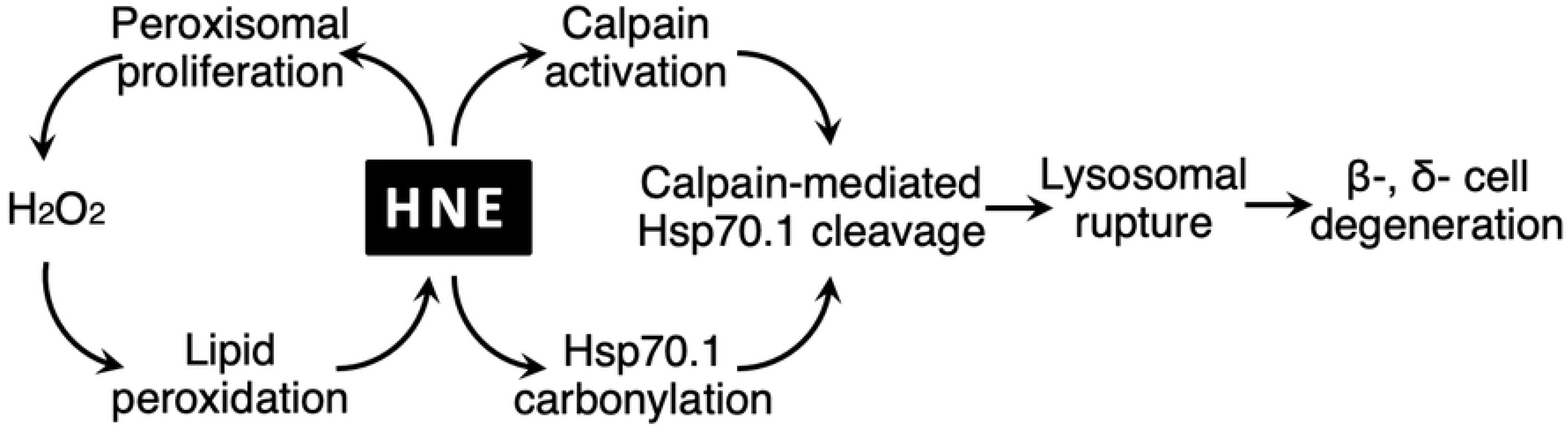
The molecular cascade explaining degeneration of β- and δ-cells in the Langerhans islet of the monkey after hydroxynonenal injections. Peroxisomes play a crucial role for the sustained H_2_O_2_ production, while calpain-mediated cleavage of carbonylated Hsp70.1 causes lysosomal membrane rupture/permeabilization. In both cascades, hydroxynonenal plays a central role for the β- and δ-cell degeneration.

Concerning the molecular mechanism of ischemic neuronal death, Yamashima et al. formulated the ‘calpain-cathepsin hypothesis’ in 1998 [14]. Thereafter, concerning the mechanism of Alzheimer’s neuronal death, they suggested that Hsp70.1 with dual functions of molecular chaperone and lysosomal stabilizer becomes vulnerable to the cleavage by activated μ-calpain, which is facilitated after Hsp70.1 carbonylation by hydroxynonenal [33,69,70,71]. Due to the hydroxynonenal-induced carbonylation with the subsequent calpain-mediated cleavage of carbonylated Hsp70.1, functional Hsp70.1 decreases steadily and this results in both accumulation of autophagosomes and permeabilization/rupture of the lysosomal membrane (Fig. 8). By inducing Hsp70.1 disorder, hydroxynonenal conceivably plays sinister roles in the occurrence of diverse lifestyle diseases [11].

## Conclusions

By oxidizing Hsp70.1, both the dietary ω-6 fatty acid-(exogenous) and the biomembrane phospholipid-(intrinsic) peroxidation product ‘hydroxynonenal’, when combined, conceivably play crucial roles for the occurrence of the Langerhans cell degeneration. The authors speculate from the present monkey experimental paradigm that ‘hydroxynonenal’ might be a real culprit behind type 2 diabetes.

## LIST OF ABBREVIATIONS

ER: endoplasmic reticulum
Hsp70.1: heat-shock protein 70.1
H_2_O_2_: hydrogen peroxide
OH^•^: hydroxyl radicals
NEFAs: nonesterified fatty acids
ROS: reactive oxygen species
O_2_^•−^: superoxide radicals

## Acknowledgements

The authors are deeply indebted to Mr. Jun Uchimoto for the daily care of monkeys and autopsy assistance, Mrs. Katsumi Hara and Mrs. Masayo Baba for the tissue preparation, and Mrs. Rie Nishioka and Mrs. Mai Nakayama for the secretory work.

